# Apoptosis-induced FGF signalling promotes non-cell autonomous resistance to cell death

**DOI:** 10.1101/2020.07.12.199430

**Authors:** Florian J. Bock, Catherine Cloix, Desiree Zerbst, Stephen W.G. Tait

## Abstract

Damaged or superfluous cells are often eliminated by apoptosis. Although a cell-autonomous process, apoptotic cells communicate with their environment in different ways. However, the extent to which apoptotic cells alerting their neighbours to potential danger is unclear. Addressing this question, here we describe a mechanism whereby dying cells can promote survival of neighbouring cells. We find that during apoptosis, cells release the growth factor FGF2, leading to MEK/ERK-dependent transcriptional upregulation of pro-survival BCL-2 proteins in a non-cell autonomous manner. This transient upregulation of prosurvival BCL-2 proteins in turn can protect neighbouring cells from apoptosis. Accordingly, we find in certain cancer types a correlation between FGF-signalling, BCL-2 expression and worse prognosis. Importantly, either co-treatment with FGF-receptor inhibitors or removal of apoptotic stress restores apoptotic sensitivity. These data reveal a pathway by which dying cells can increase resistance to cell death in surrounding cells. Beyond mediating cytotoxic drug resistance, this process may serve additional roles, for instance limiting tissue damage in response to stress.

## Introduction

The cellular decision to live or die is of fundamental importance across biology. For instance, inappropriate cell survival has been linked to various diseases including cancer and autoimmunity ^1^. In cancer, many therapies act by engaging apoptosis, and the degree of apoptotic sensitivity or “apoptotic priming” often correlates with therapeutic efficacy ^1,2^. Because of this, understanding how cancer cells survive therapy should provide new ways to circumvent this and improve tumour cell killing.

Mitochondrial apoptosis represents a major form of regulated cell death ^3^. During apoptosis, mitochondria are permeabilised through a process called mitochondrial outer membrane permeabilisation or MOMP. Widespread MOMP effectively acts as cellular death sentence by releasing mitochondrial proteins, such as cytochrome *c*, that activate caspase proteases leading to rapid apoptosis ^3^. Even in the absence of caspases, MOMP typically commits a cell to death, and is thus considered a point-of-no-return. Consequently, mitochondrial outer membrane integrity is tightly regulated by pro- and anti-apoptotic BCL-2 family proteins. A new class of anti-cancer drugs, called BH3-mimetics, target anti-apoptotic BCL-2 proteins, thereby sensitising cancer cells to apoptosis ^4^. The BCL-2 specific BH3-mimetic venetoclax ^5^ shows considerable clinical promise and is approved for treatment in chronic lympocytic leuakaemia (CLL) ^6^ and as combination therapy in acute myeloid leukaemia (AML) ^7,8^. However, in solid tumours, BH3-mimetics are typically less effective, suggesting additional survival mechanisms must be targeted in order to maximise their potential.

In this study, we set out to identify mechanisms of apoptotic resistance using BH3-mimetics as tool compounds. Selecting for cells surviving venetoclax treatment, we found that resistance was associated with increased anti-apoptotic BCL-2 and MCL1 expression. Surprisingly, resistance occurred in a non-cell autonomous manner. We find that as cells undergo apoptosis, they can release FGF2. In turn FGF2 triggers MEK/ERK signalling, resulting in increased anti-apoptotic BCL-2 and MCL1 protein expression and apoptotic resistance. In certain cancer types, we found a correlation between FGF-signalling, BCL-2 and MCL1 expression and poorer patient prognosis, suggesting its relevance in vivo. These findings unveil an unexpected non-cell autonomous mechanism of apoptotic resistance, where cell death – via FGF signalling – promotes cell survival. As we discuss, this process may have wide-ranging roles in health and disease.

## Results

### BH3-mimetics and BH3-only proteins upregulate BCL-2 and MCL1 causing apoptotic resistance

Our initial goal was to define mechanisms of cell death resistance using BCL-2 targeting BH3-mimetics as tool compounds. For this purpose, we used our recently developed method called mito-priming ^9^. In this system, cells co-express a pro-apoptotic BH3-only protein and an anti-apoptotic BCL-2 family member at equimolar levels and are therefore highly sensitive to BCL-2 targeting BH3-mimetic drugs (**Figure 1a**). HeLa cells were used that stably express tBID together with BCL-2 (HeLa tBID-2A-BCL-2, hereafter called HeLa tBID2A). Cell viability was then determined following venetoclax (ABT-199) treatment using Sytox green exclusion and Incucyte live-cell imaging. As expected, the majority of cells died rapidly following venetoclax treatment, nevertheless some cells failed to die (**Figure 1b**). To investigate the mechanism of venetoclax resistance, HeLa tBID2A cells were cultured continuously in venetoclax to select for resistant cells. Increased expression of pro-survival BCL-2 family proteins is a common means of apoptotic resistance ^10^. Indeed, cells that were continuously or intermittently cultured in venetoclax displayed higher expression of anti-apoptotic BCL-2 and MCL1 (**Supplementary Figure 1a**). Surprisingly, following culture in regular medium post-venetoclax treatment, the resistant cells became sensitive to venetoclax again over time (**Supplementary Figure 1b**). This decrease of resistance was accompanied by a decrease in BCL-2 and MCL1 expression back to basal levels (data not shown). We next investigated whether short term treatment with venetoclax was sufficient to promote BCL-2 and MCL1 upregulation. Indeed, 3 hours of venetoclax treatment led to increased levels of BCL-2 and MCL1 (**Figure 1c**). As before, BCL-2 and MCL1 levels decreased following removal of venetoclax, demonstrating a reversible upregulation (**Figure 1d**). This effect was not restricted to venetoclax, because an increase in BCL-2 and MCL1 expression was also observed following treatment with other BH3 mimetics (ABT-263 and ABT-737) (**Supplementary Figure 1c).** Because venetoclax induced upregulation of BCL-2 and MCL1 is rapidly reversible this suggests that it is not genetics based. We noted an initial resistance of HeLa tBID2A cells cultured in venetoclax to re-treatment with venetoclax (**Supplementary Figure 1b**), presumably due to increased levels of BCL-2 and MCL1. Because increased levels of BCL-2 and MCL1 were also observed in response to acute treatment, we investigated if this was also sufficient to protect from apoptosis. Indeed, treatment with venetoclax for 48 hours could completely protect from re-treatment with venetoclax and S63845, a specific inhibitor of MCL1 ^11^ (**Figure 1e**). This protection was dependent on the increased levels of BCL-2 and MCL1, because increasing the dose of venetoclax and S63845 could overcome the resistance (**Figure 1e**). The pro-apoptotic proteins BAX and BAK are essential for mitochondrial outer membrane permeabilization (MOMP) during apoptosis ^12^. To determine the role of apoptosis in the upregulation of BCL-2 and MCL1, we generated HeLa tBID2A cells deficient in BAX and BAK using CRISPR/Cas9 genome editing. As expected, BAX/BAK-deleted HeLa tBID2A cells were completely protected from mitochondrial apoptosis and caspase activation in response to venetoclax treatment (**Supplementary Figure 1d, e**). Nevertheless, despite the inability to undergo apoptosis, BAX/BAK-deleted HeLa tBID2A cells still upregulated BCL-2 and MCL1 (**Figure 1f, Supplementary Figure 1f**). Similarly, upregulation of BCL-2 and MCL1 was also observed when caspase activity was blocked using the caspase inhibitor qVD-OPh (**Figure 1f).** These data demonstrate that while the upregulation of BCL-2 and MCL1 requires BH3-mimetic treatment, it occurs irrespective of apoptosis. Finally, to determine whether upregulation of BCL-2 and MCL1 was specific to BH3-mimetics, we examined if a comparable effect could be observed by expressing BH3-only proteins. Control or BAX and BAK deleted HeLa cells were transfected with BH3-only proteins (tBID, PUMA, tBID (BIM BH3)) and analysed for MCL1 expression by western blot (**Figure 1g, Supplementary Figure 1g**). In all cases, MCL1 expression was upregulated, indicating that BH3-only proteins can fulfil a similar function as BH3 mimetics. Collectively, these data demonstrate that BH3-mimetics and BH3-only proteins can promote apoptotic resistance by increasing pro-survival BCL-2 protein expression.

**Figure 1:**
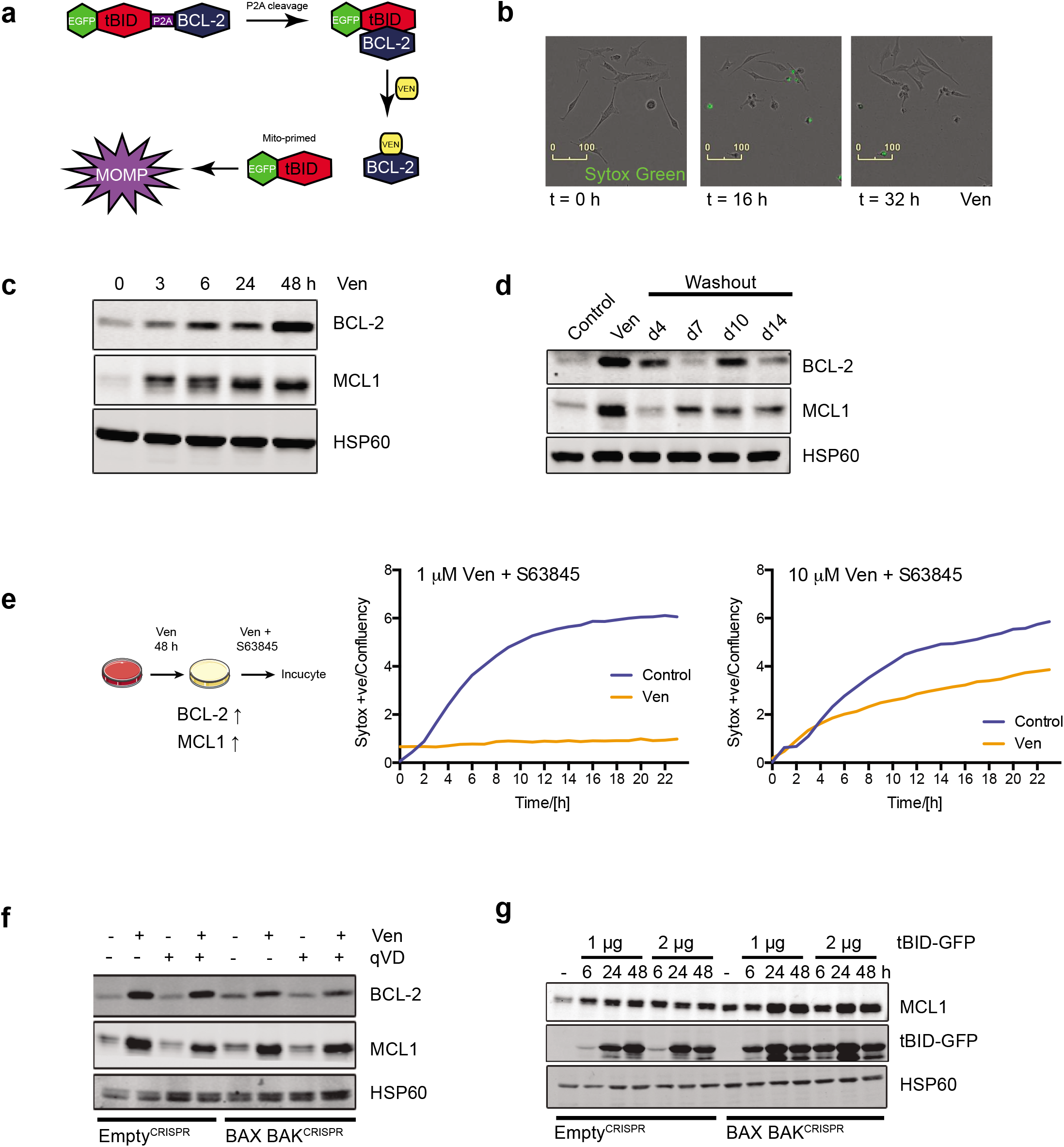
BH3-mimetics and BH3-only proteins upregulate BCL-2 and MCL1 causing apoptotic resistance. (a) Schematic model of the mitoprimed system (b) HeLa tBID2A cells were treated with venetoclax and imaged over time. Dead cells were stained with Sytox green. Selected image stills are shown. Scale bar: 100 μm, representative images of > 3 independent experiments (c) HeLa tBID2A cells were treated with 500 nM venetoclax, harvested at the indicated time points and protein expression was analysed by western blot. Representative images from two independent experiments (d) HeLa tBID2A cells were treated for 48 h with 500 nM venetoclax followed by replacement with regular growth medium (washout). At the indicated times post medium change cells were harvested and protein expression was analysed by western blot. Representative images from two independent experiments (e) HeLa tBID2A cells were treated with or without venetoclax as indicated for 48 h followed by treatment with venetoclax and S63845. Cell death was then monitored by Sytox Green staining and Incucyte imaging. Representative data from two independent experiments (f) Control or BAX BAK^CRISPR^ HeLa tBID2A cells were treated with 500 nM venetoclax in combination with 10 μM qVD-oph as indicated for 48 h, harvested and protein expression was analysed by western blot. Representative images from two independent experiments (g) Control or BAX BAK^CRISPR^ HeLa cells were transfected with the indicated amounts of tBID-GFP plasmid, harvested after the indicated times and protein expression was analysed by western blot. Representative images from two independent experiments

### BH3-mimetics and BH3-only proteins can upregulate anti-apoptotic BCL-2 proteins in a non-cell autonomous manner

To understand how venetoclax treatment causes upregulation of BCL-2 and MCL1, we determined whether increases in protein or mRNA stability might contribute. HeLa tBID2A cells were treated with venetoclax for 24 hours, after which inhibitors of protein synthesis (cycloheximide) or transcription (actinomycin D) were added for varying times. Both MCL1 and BCL-2 protein or mRNA stability was similar after venetoclax treatment (**Supplementary Figures 2a, b**), indicating that mechanisms besides protein or mRNA stability are likely responsible. Given this, we investigated if venetoclax might upregulate BCL-2 and MCL1 in a non-cell autonomous manner. Control or BAX/BAK deleted HeLa tBID2A cells were treated for 3 hours with 500 nM venetoclax, followed by exchange to regular medium for 45 hours. Media from treated cells was then transferred to recipient cells, which were examined for BCL-2 and MCL1 expression after 48 hours (**Figure 2a**). Importantly, media from venetoclax treated HeLa tBID2A cells promoted upregulation of BCL-2 and MCL1 in recipient cells (**Figure 2b**). Similarly, supernatant from BAX and BAK deficient cells also promoted MCL-1 and BCL-2 upregulation, supporting earlier findings that cell death is not required for this effect (**Figure 2b**). Supernatant from venetoclax treated cells failed to induce apoptosis in recipient cells, demonstrating absence of potentially residual venetoclax (**Supplementary Figure 2c**). Additionally, media from HeLa tBID2A cells treated with a different BCL-2 inhibitor, S55746 ^13^, also upregulated MCL1 and BCL-2 in recipient cells (**Supplementary Figure 2d**). To investigate the mechanism of this non-cell autonomous effect, we first characterised the signal causing upregulation of anti-apoptotic BCL-2 proteins. Supernatant from venetoclax treated HeLa tBID2A cells was subjected to centrifugal filtration using a filter with a 3 kDa cut-off. Flow-through and concentrate was added to recipient cells for 48 hours, and MCL1 and BCL-2 expression was determined by western blot (**Supplementary Figure 2e**). Only the concentrate (containing molecules above 3 kDa) was capable of increasing BCL2 and MCL-1 expression, suggesting that small molecules such as metabolites and lipids are not responsible. Importantly, Proteinase K treatment of supernatant from BH3-mimetic treated cells abolished the ability to upregulate MCL1 and BCL-2, consistent with the factor(s) being of protein nature (**Figure 2c**). Finally, we investigated whether BH3-only proteins can also have a similar non-cell autonomous effect. HeLa or 293T cells were transfected with tBID and the supernatant was transferred onto recipient cells. Consistent with earlier results, supernatant transferred from tBID transfected cells also caused an up-regulation in BCL-2 and MCL1 expression (**Figure 2d, Supplementary Figure 2f**). Together, these data demonstrate that BH3-mimetics and BH3-only proteins can upregulate BCL-2 and MCL1 expression in a non-cell autonomous manner.

**Figure 2:**
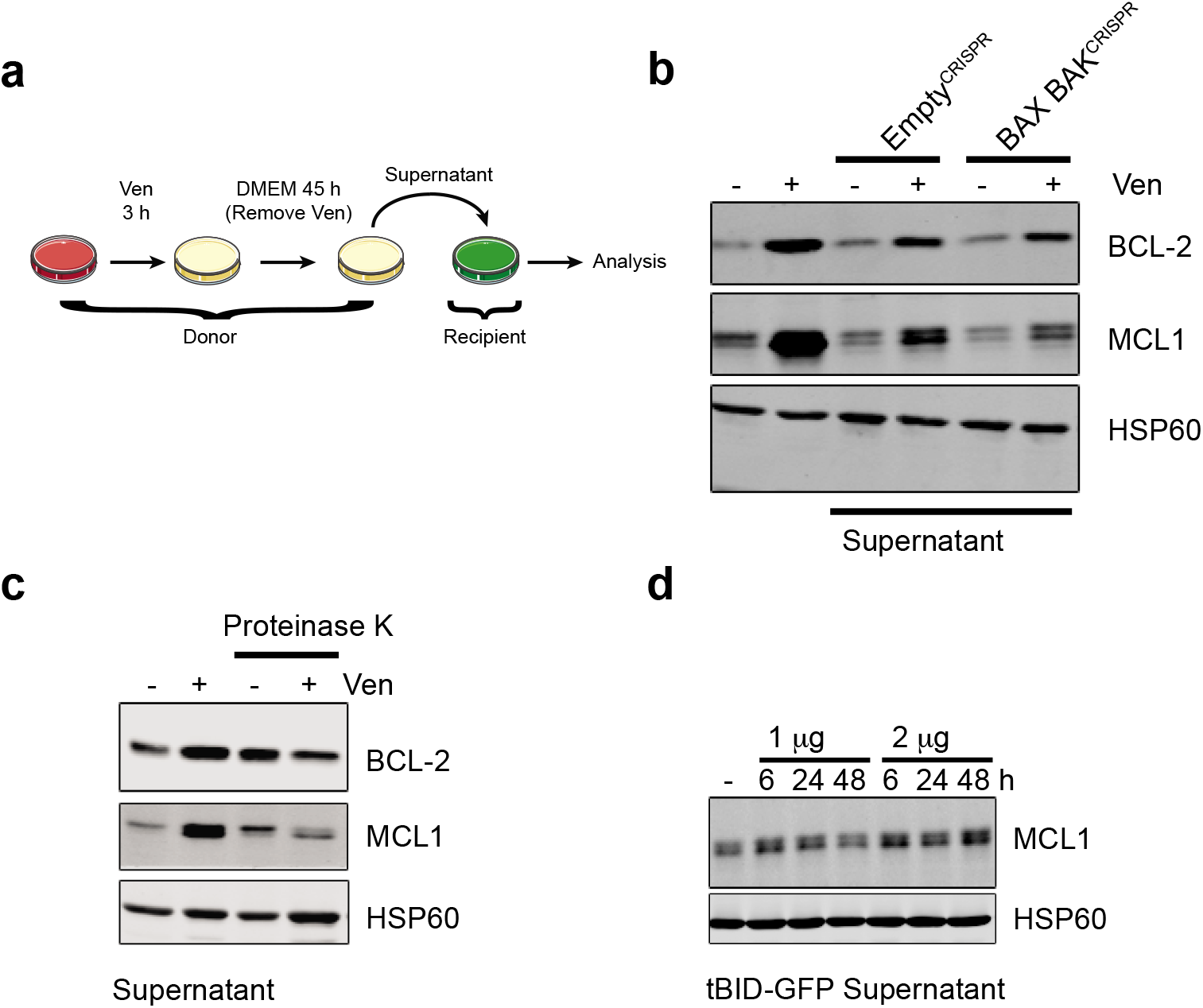
BH3-mimetics and BH3-only proteins can upregulate anti-apoptotic BCL-2 proteins in a non-cell autonomous manner. (a) Schematic of supernatant transfer experiments: HeLa tBID2A cells were treated with 500 nM venetoclax for 3 h, followed by exchange to regular growth medium for 45 h. Supernatant was harvested, filtered and added onto recipient cells before analysis (b) Supernatant from untreated or venetoclax treated control or BAX BAK^CRISPR^ HeLa tBID2A cells was harvested as described in (a) and added onto control HeLa tBID2A cells for 48 h followed by harvesting and analysis of protein expression by western blot. Representative images from three independent experiments (c) Supernatant from control or venetoclax treated cells was digested with Proteinase K before addition onto recipient cells and protein expression was analysed by western blot after 48 h. Representative images from two independent experiments (d) Supernatant from HeLa cells transfected with tBID-GFP was harvested after the indicated times and transferred onto recipient HeLa cells. After 48 h, recipient cells were harvested and protein expression analysed by western blot. Representative images from two independent experiments

### Non-cell autonomous upregulation of anti-apoptotic BCL-2 proteins requires MEK/ERK signalling

We next sought to define the non-cell autonomous mechanism causing anti-apoptotic BCL-2 protein upregulation. For this purpose, HeLa tBID2A cells were treated with venetoclax together with inhibitors targeting pathways previously implicated in anti-apoptotic BCL-2 regulation ^14–17^. After co-treatment for 48 hours, cell lysates were probed for BCL-2 and MCL1 expression by western blot. Of all the tested inhibitors, only trametinib (a MEK kinase inhibitor ^18^) potently blocked venetoclax induced BCL-2 and MCL1 expression (**Figures 3a, 3b**). The decrease in phosphorylation of ERK1/2, a direct target of MEK ^19^, validated trametinib activity (**Figure 3b**). Upregulation of BCL-2 and MCL1 was transcriptional, because venetoclax treatment increased RNA levels, which could be inhibited by trametinib co-treatment (**Figure 3c)**. Upregulation of MCL1 by BH3 mimetics was not limited to HeLa cells, since it was also observed in IMR90 and CWR-R1 cells (**Supplementary Figures 3a and 3b).** We next investigated whether non-cell autonomous upregulation of BCL-2 and MCL1 was also dependent on MEK/ERK signalling. Trametinib was added to supernatant from venetoclax treated HeLa tBID2A or CWR-R1 cells before the supernatant was added to recipient cells. BCL-2 and MCL1 upregulation was effectively blocked by trametinib (**Figure 3d, Supplementary Figure 3c**). Again this effect was transcriptional, since upregulation of MCL1 RNA in recipient cells was inhibited by trametinib addition to the supernatant (**Supplementary Figure 3d**). Furthermore, supernatant of venetoclax treated cells could directly stimulate MEK activity in recipient cells as determined by increased p-ERK levels (**Figure 3e**). To further investigate these findings, we generated ERK1/2 deficient HeLa tBID2A cells by CRISPR/Cas9 genome editing (**Figure 3f**). Media from control or ERK1/2^CRISPR^ venetoclax treated cells was transferred onto control or ERK1/2^CRISPR^ cells. After 48 hours, BCL2 and MCL-1 expression was determined by western blot. BCL-2 and MCL1 expression increased following incubation with media from venetoclax treated cells or after direct treatment with venetoclax in control cells, but was severely attenuated in ERK1/2^CRISPR^ cells (**Figure 3f**). Finally, we investigated whether MEK signalling, by enabling BCL-2 and MCL1 upregulation, contributed to venetoclax resistance. HeLa tBID2A cells were incubated with venetoclax ± trametinib for 48 hours, after which they were treated with venetoclax and S63845 and cell viability was determined by Sytox Green exclusion and Incuyte live cell imaging (**Supplementary Figure 3e**). Whereas venetoclax pre-treated cells were resistant, trametinib co-treatment completely abolished this resistance, supporting a functional role for MEK signalling in mediating apoptotic resistance via BCL-2 and MCL1 upregulation. Similarly, transfer of venetoclax treated supernatant conferred resistance of the recipient cells. The resistance was dependent on BCL-2 and MCL1 upregulation, because supplementing the venetoclax treated supernatant with trametinib before addition to recipient cells re-sensitised those cells to the cytotoxic treatment (**Figure 3g**). These data show that BH3-mimetics activate MEK-ERK signalling, causing upregulation of BCL-2 and MCL-1 and apoptotic resistance in a non-cell autonomous manner.

**Figure 3:**
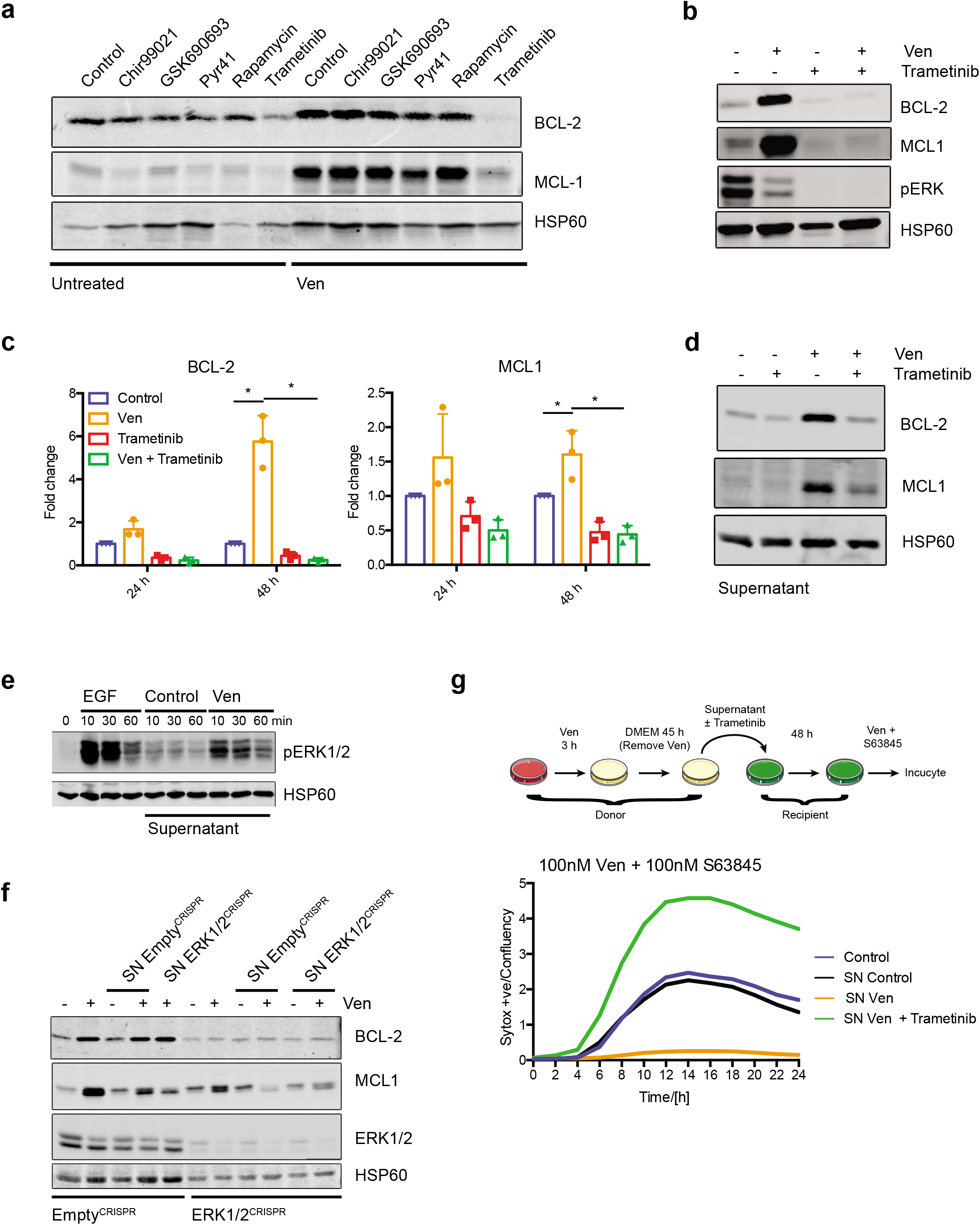
Non-cell autonomous upregulation of anti-apoptotic BCL-2 proteins requires MEK/ERK signalling. (a) HeLa tBID2A cells were untreated or treated with 500 nM venetoclax in combination with the indicated inhibitors for 48 h, harvested and protein expression was analysed by western blot. Representative images from two independent experiments (b) HeLa tBID2A cells were untreated or treated with 500 nM venetoclax in combination with 500 nM Trametinib for 48 h, harvested and protein expression was analysed by western blot. Representative images from three independent experiments (c) HeLa tBID2A cells were untreated or treated with 500 nM venetoclax in combination with 500 nM Trametinib, harvested and RNA expression was analysed by RT-qPCR. n = 3 independent experiments; mean values ± s.d., *: p < 0.05, Tukey corrected one-way ANOVA (d) Supernatant from untreated or venetoclax treated HeLa tBID2A cells was treated with 500 nM Trametinib before addition onto recipient cells. After 48 h of incubation, recipient cells were harvested and protein expression analysed by western blot. Representative images from three independent experiments (e) Supernatant from untreated or venetoclax treated HeLa tBID2A cells was added onto recipient cells. After the indicated times, recipient cells were harvested and protein expression analysed by western blot. Treatment with EGF served as a positive control. Representative images from two independent experiments (f) Control or ERK1/2^CRISPR^ HeLa tBID2A cells were treated directly with venetoclax or with supernatant from control or ERK1/2^CRISPR^ HeLa tBID2A cells as indicated before harvesting and western blot analysis for protein expression. Representative images from two independent experiments (g) Supernatant from control or venetoclax treated HeLa tBID2A cells was supplemented with 500 nM Trametinib as indicated before addition onto recipient cells. After 48 h of incubation, the recipient cells were treated with 100 nM venetoclax and 100 nM S63845 and survival was monitored by Sytox green exclusion and live cell imaging Representative data from two independent experiments

### FGF signalling mediates non-cell autonomous upregulation of BCL-2 proteins

Various ligands can bind to receptors that signal through MEK/ERK, with receptor tyrosine kinases (RTK) being prominent activators of this pathway. Therefore, to determine the paracrine mediator(s) causing BCL-2 and MCL1 upregulation, we initially focussed on the RTK pathway. To identify potential ligands present after BH3-mimetic treatment, supernatant from control or venetoclax treated HeLa tBID2A cells was incubated with a human growth factor antibody array enabling detection of 41 different growth factors (**Figure 4a**). One of the ligands with increased levels in venetoclax treated supernatant was FGF2, which was corroborated by subsequent ELISA analysis (**Figure 4b**). Addition of recombinant FGF2 to cells at a concentration similar to the one measured by ELISA in venetoclax treated supernatant was sufficient to upregulate BCL-2 and MCL1 expression (**Figure 4c**). To directly address the importance of FGF2 in mediating paracrine upregulation of BCL-2 proteins, we generated FGF2 deficient cell lines by CRIPSR/Cas9. Loss of FGF2 completely supressed the ability of venetoclax to upregulate BCL-2 and MCL1 in a paracrine manner, supporting a key role for FGF2 (**Figure 4d**). Consistent with activation of FGF-signalling, known target genes of FGF receptors (CDX2, DUSP6 and SPRY4) were also upregulated in response to supernatant from venetoclax treated cells (**Figure 4e**). This upregulation could be blocked by adding trametinib to the supernatant before addition to recipient cells. To investigate the requirement of FGF receptors for upregulation of BCL-2 and MCL1, we used two different FGFR inhibitors (AZD4547 and PRN1371 ^20,21^). Co-treatment of HeLa tBID2A cells with either inhibitor and venetoclax prevented BCL-2 and MCL1 upregulation (**Supplementary Figure 4a**). Similarly, supernatant from venetoclax treated HeLa tBID2A cells supplemented with FGFR inhibitors prevented upregulation of BCL-2 and MCL1 on recipient cells (**Figure 4f**). Reduced upregulation of MCL1 was also observed upon co-treatment of MRC-5 cells with venetoclax and PRN1371 (**Supplementary Figure 4b**). The FGF receptor family is composed of four different receptors ^22^. To determine which receptors were responsible to signal the upregulation of BCL-2 and MCL1 by FGF2, we used RNAi to knockdown their expression individually or in combination. Knocking down FGFR1 and FGFR3, either individually or in combination, prevented upregulation of BCL-2 and MCL1 either after direct treatment (**Figure 4g)** or with venetoclax treated supernatant (**Figure 4h**). In contrast, knockdown of FGFR2 and FGFR4 failed to affect BCL-2 and MCL1 expression (**Supplementary Figures 4c - f**). Collectively, these data demonstrate that FGF-signalling, triggered by release of FGF-2 from apoptotic cells, is required and sufficient for non-cell autonomous upregulation of anti-apoptotic BCL-2 proteins.

**Figure 4:**
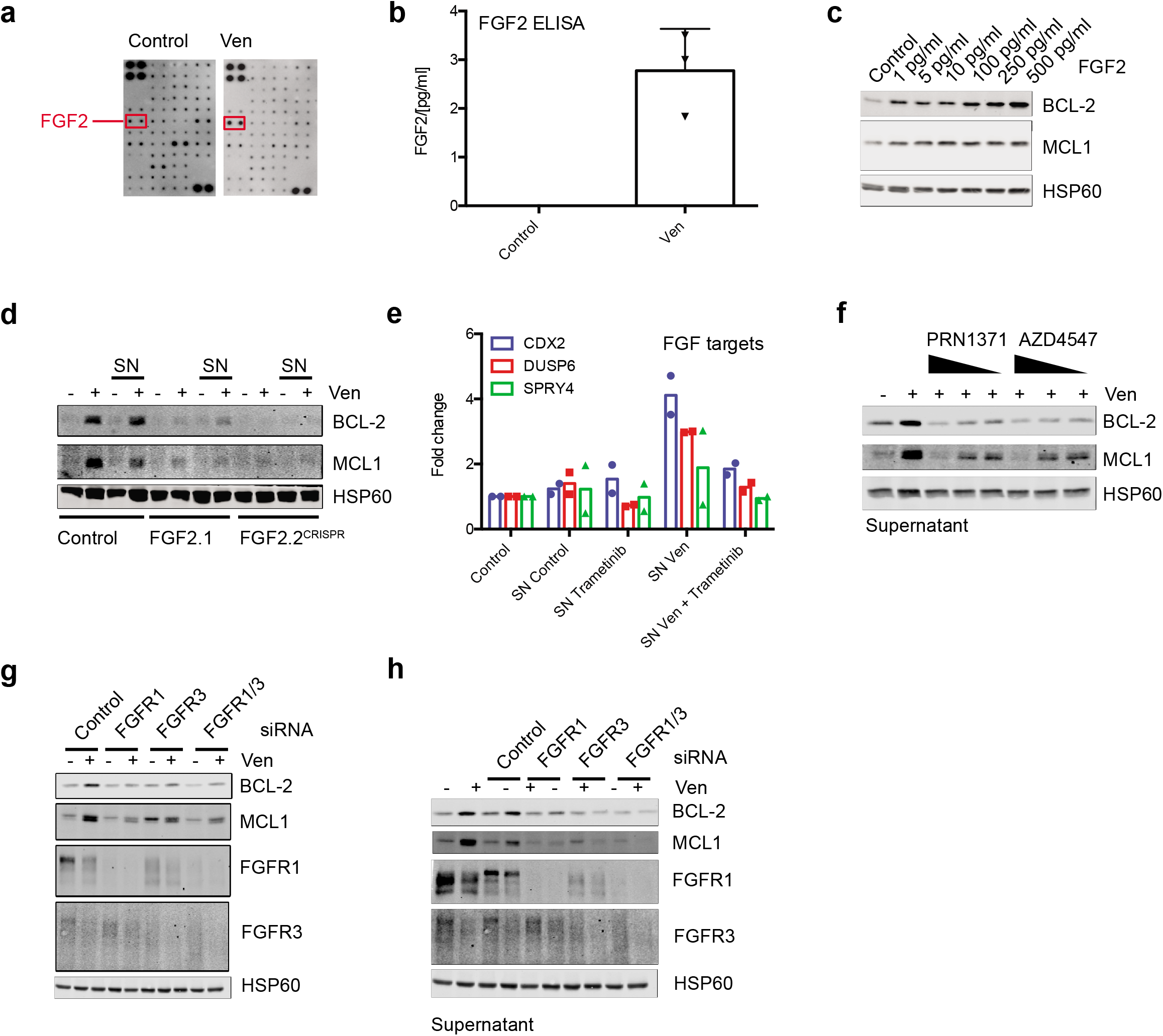
FGF signalling mediates non-cell autonomous upregulation of BCL-2 proteins. (a) A growth factor membrane ligand array was probed with supernatant from control or venetoclax treated HeLa tBID2A cells. Spots for FGF2 are indicated. Representative images from two independent experiments (b) Levels of FGF2 were determined in supernatant from control or venetoclax treated HeLa tBID2A cells by ELISA. Representative data from three independent experiments (c) HeLa tBID2A cells were treated with recombinant FGF2, harvested after 6 h and protein expression analysed by western blot. Representative images from three independent experiments (d) Control or FGF2^CRISPR^ HeLa tBID2A cells were directly treated with venetoclax or the respective supernatant for 48 h as indicated before harvesting and analysis of protein expression by western blot. Representative images from two independent experiments (e) Supernatant from untreated or venetoclax treated HeLa tBID2A cells was treated with 500 nM Trametinib before addition onto recipient cells. After 3 h, recipient cells were harvested and RNA expression of FGF target genes analysed by RT-qPCR. Representative data from two independent experiments (f) Supernatant from control or venetoclax treated HeLa tBID2A cells was treated with decreasing doses of FGFR inhibitors as indicated before addition onto recipient cells for 48 h and analysis for protein expression by western blot. Representative images from three independent experiments (g) HeLa tBID2A cells were transfected with siRNA targeting FGFR1 and FGFR3 alone or in combination for 24 h before addition of 500 nM venetoclax, harvesting after 48 h and analysis of protein expression by western blot. Representative images from three independent experiments (h) HeLa tBID2A cells were transfected with siRNA targeting FGFR1 and FGFR3 alone or in combination for 24 h before addition of control or venetoclax treated supernatant from control cells, harvesting after 48 h and analysis of protein expression by western blot. Representative images from three independent experiments

### FGF signalling is essential for non-cell autonomous apoptotic resistance

We next investigated the contribution of FGF mediated BCL-2 and MCL1 upregulation to apoptosis resistance. First, HeLa tBID2A cells were co-treated with venetoclax and inhibitors of FGF signalling (PRN1371, AZD4547) or MEK (trametinib). Whereas the venetoclax treated cells were resistant to apoptosis induced by retreatment with venetoclax and S63845, cells co-treated with FGF signalling inhibitors were re-sensitized to apoptosis (**Figure 5a**). As observed previously, increasing the dose of venetoclax and S63845 could restore sensitivity to venetoclax pretreated cells (**Supplementary Figure 5a**). Next, the potential of FGF signalling to promote non-cell autonomous apoptotic resistance was investigated. Supernatant was harvested from venetoclax treated HeLa tBID2A cells and supplemented with inhibitors of the FGF signalling pathway before addition onto recipient cells. Consistent with our earlier data, supernatant from venetoclax treated cells conferred apoptotic resistance to recipient cells (**Figure 5b**). Crucially, apoptotic sensitivity was restored upon inhibition of either FGF or MEK signalling. Again, increasing the dose of venetoclax and S63845 could kill the venetoclax pre-treated resistant cells (**Supplementary Figure 5b**). To investigate this further, we assessed effects on long-term clonogenic survival. Supporting earlier data, supernatant from venetoclax treated cells could confer long-term protection from apoptosis, which was restored by supplementation of FGF signalling inhibitors (**Figure 5c**). These data support a model whereby following apoptotic stress, cells can signal non-cell autonomous apoptotic resistance by FGF-signalling dependent on the upregulation of BCL-2 and MCL1 (**Figure 5d**). Finally, we investigated whether a similar mechanism may promote cancer. Interrogating the TCGA, we first determined an FGF pathway activation score for specific cancer types by examining FGF target gene expression. From this, we examined MCL1 and BCL-2 expression and determined correlation to FGF target gene expression. Using this approach, we identified several cancer types that displayed a correlation between FGFR activation and BCL-2 or MCL1 expression (**Figure 5e, Supplementary Figures 6 a, b**). Next, we determined whether this correlation had an influence on disease progression by separating samples into groups based on FGF score and BCL-2 or MCL1 expression. Strikingly, in some but not all cancer types, survival of the high scoring group was significantly decreased compared to the low scoring group (**Figure 5e, Supplementary Figure 6a)**. This suggests that FGF mediated resistance might have a protective effect on cancer cell survival and therefore worsen prognosis.

**Figure 5:**
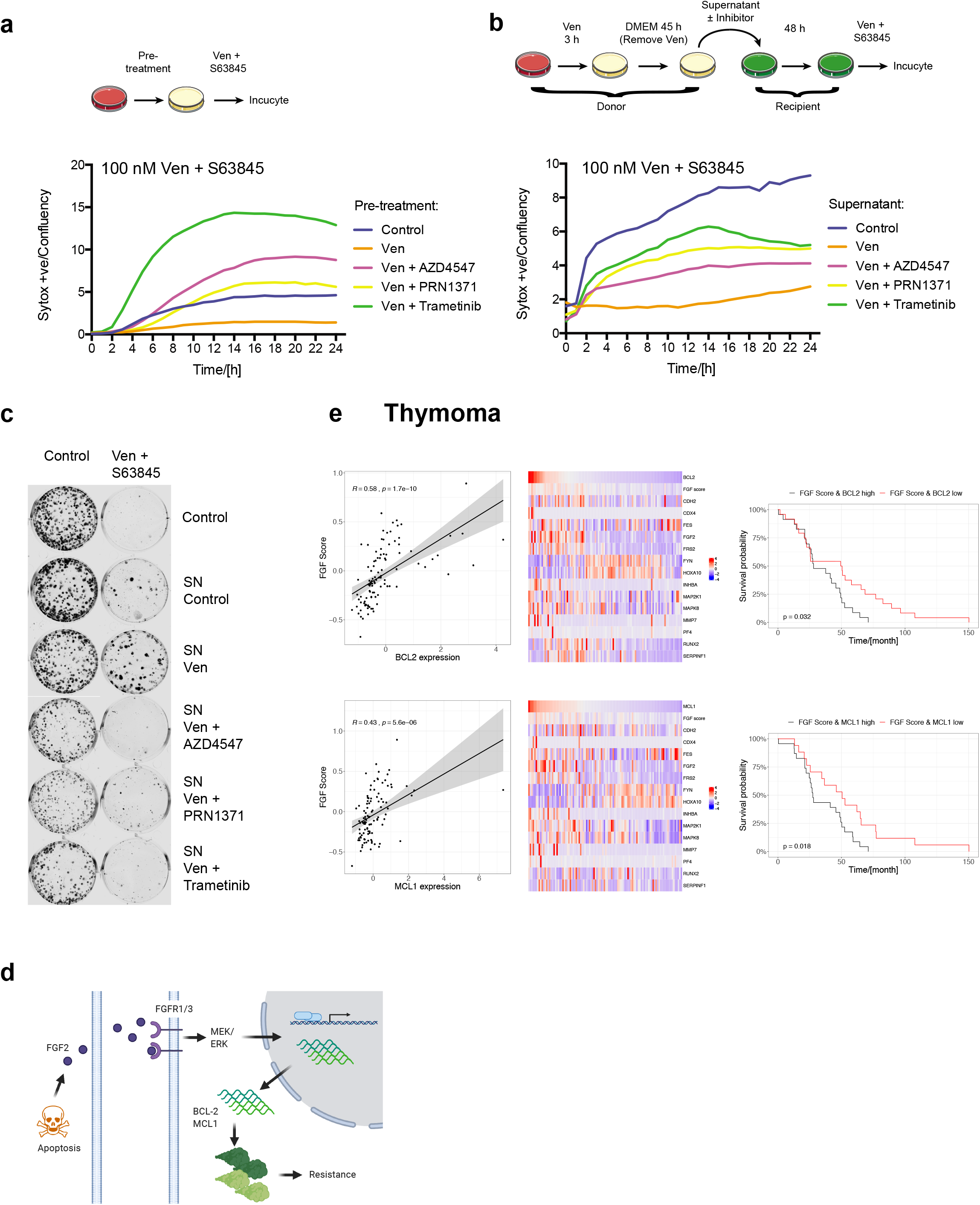
FGF signalling is essential for non-cell autonomous apoptotic resistance. (a) HeLa tBID2A cells were treated with 500 nM venetoclax in combination with RTK pathway inhibitors (AZD4547 (5 μM), PRN1371 (10 μM) or Trametinib (500 nM)) as indicated for 48 h. Then the cells were treated with 500 nM venetoclax + S63845 and cell survival was monitored by Incucyte. Representative data from two independent experiments (b) HeLa tBID2A cells were incubated with 500 nM venetoclax treated supernatant supplemented with RTK pathway inhibitors (AZD4547 (5 μM), PRN1371 (10 μM) or Trametinib (500 nM)) as indicated before addition onto recipient cells for 48h. Then the cells were treated with 100 nM venetoclax + S63845 and cell survival was monitored by Incucyte. Representative data from two independent experiments (c) HeLa tBID2A cells were plated at low density (1000 cells per 6 well) and incubated with 500 nM venetoclax treated supernatant supplemented with RTK pathway inhibitors (AZD4547 (5 μM), PRN1371 (10 μM) or Trametinib (500 nM)) as indicated for 48 h. Cells were then treated with 2.5 μM venetoclax + S63845 for another 48 h before replacement with regular growth medium. After an additional 5 days colonies were visualised by crystal violet staining. Representative images from two independent experiments (d) Working model (e) Correlation between FGF Score and BCL-2 (upper left) or MCL1 (lower left) in the TCGA Thymoma dataset. FGF score, FGF receptor target gene and BCL-2 (upper middle) or MCL1 (lower middle) expression in the TCGA Thymoma dataset. Survival of TCGA Thymoma patients stratified by high FGF score and BCL-2 (upper right) and MCL1 (lower right) expression

## Discussion

Innate or acquired resistance to cell death is fundamentally important in health and disease. In cancer, evasion of apoptosis can both promote cancer and inhibit treatment response, leading to tumour relapse ^23^. To understand how cells can resist cell death, in this study we used BCL-2 targeting BH3-mimetics as tool compounds. Unexpectedly, we uncovered a non-cell autonomous mechanism that enables apoptotic resistance. We found that during apoptosis, cells can release the growth factor FGF2. By activating MEK/ERK signalling, FGF2 upregulates anti-apoptotic BCL-2 protein expression ^24,25^ in neighbouring cells, protecting against apoptosis. Importantly, resistance can be overcome by co-treatment of BH3-mimetics with FGFR-inhibitors, demonstrating the functional significance of the pathway. Finally, we describe a correlation between increased FGF signalling, anti-apoptotic BCL2 protein expression and poor patient prognosis, suggesting its relevance *in vivo*. As we discuss further, the process we describe here, where cell death promotes life, may have various pathophysiological functions.

Most cancer therapies work by killing tumour cells, consequently resistance to cell death profoundly impacts on therapeutic efficacy ^23^. Typically, cancer cells can evade apoptosis through inactivating mutations in pathways that sense damage or activate cell death ^23^. For instance, in chronic lymphocytic leukemia (CLL), BCL-2 mutations have recently been reported that render BCL-2 unable to effectively bind the BH3-mimetic venetoclax, causing apoptotic resistance ^26–29^. The resistance mechanism we report here is not mutation based, but instead is due to dying cells releasing FGF2 that causes transient apoptotic resistance in surrounding cells. This mechanism fits the concept of persistence, which has emerged to explain transient, non-heritable resistance of tumour cells to therapy ^30^. Cancer cells that transiently evade cell death, called persister cells, blunt the effectiveness of chemotherapy and provide a cell pool from which drug-resistant tumours may arise. The mechanism we describe herein represents a non-cell autonomous way of generating persistence (via FGF2 release) that requires activation of the core apoptotic pathway. Suggestive of its relevance *in vivo*, we find that in certain cancer types there is a correlation between FGF-signalling, anti-apoptotic BCL-2 protein expression and poor prognosis. Importantly, we find that inhibition of FGF receptor signalling or of downstream MEK/ERK signalling greatly impedes the ability of apoptotic cells to promote survival in a non-cell autonomous manner. Our data show that apoptotic resistance is transient lasting multiple days, after which cells become sensitised again to BH3-mimetics. Extrapolating these findings to a clinical setting, this suggest the possibility that intermittent dosing, employing so-called “drug holidays”, may minimise apoptotic resistance.

The process we describe in this work – where dying cells promote survival of surrounding ones – shares characteristics with cell competition. In cell competition, cells deemed unfit (loser cells) are eliminated and can promote the survival of surrounding, winner cells ^31^. Applying this concept to our data, stressed or unfit cells within a tissue or tumour might not only be eliminated, but even increase the resistance of their neighbour cells to apoptotic stress. In general, the pathway described in this work can act as a way of tissue protection to limit damage subjected upon a tissue, where a stressed cell alerts its surrounding environment to increase the threshold of programmed cell death.

A remaining question is how apoptotic stress leads to FGF2 release in dying cells. Importantly, while FGF2 release occurs during caspase-dependent apoptosis, it is not dependent on apoptosis - neither inhibition of caspase function nor MOMP prevents FGF2 release. This suggests that neutralisation of anti-apoptotic BCL-2 proteins exerts a non-lethal signalling function, leading to FGF2 release. Indeed, a variety of non-apoptotic functions for BCL-2 have been reported previously, for instance roles in calcium signalling or metabolism ^32^. Alternatively, BH3-only proteins may be responsible for this FGF2 release. FGF2 is secreted in a non-canonical manner, which involves its binding to PI(4,5)P_2_ at the plasma membrane followed by insertion of FGF2 oligomers through the membrane ^33^. Although tBID was described to be able to interact with various membranes like artificial liposomes ^34^, lysosomes ^35^ or mitochondria ^36^, it remains to be determined if this function of tBID can be extended to FGF2 secretion.

Our data further emphasise that cell death exerts a plethora of non-cell autonomous effects. These include context dependent pro-proliferative, inflammatory and apoptotic effects ^37^. The mechanism we describe here represents an FGF-driven pro-survival effect of cell death. As discussed, this effect may have important implications for dictating therapeutic efficacy of apoptosis-inducing cancer therapy. Beyond this, cell death induced survival may have various additional effects including, but not limited to, limiting tissue damage in response to wounding or infection.

## Materials and Methods

### Cell lines and chemicals

HeLa, HeLa tBID2A BCL-2 cells ^9^, IMR90, MRC5 and 293T cells were cultured in DMEM high-glucose medium supplemented with 10% FCS and 2 mM glutamine. CWR-R1 cells were cultured in RPMI high-glucose medium supplemented with 10% FCS and 2 mM glutamine. MRC5 and IMR90 cells were cultured in 3 % oxygen. To select for venetoclax resistant cells, HeLa tBID2A BCL-2 cells were cultured continuously in the indicated dose of venetoclax for 14 days or cultured for 8 hours in venetoclax followed by 16 hours normal medium daily. The following drugs and chemicals were used: ABT-199/venetoclax (AdooQ BioScience, A12500-50), ABT-263 (ApexBio, A3007), ABT-737 (ApexBio, A8193), Actinomycin D (Calbiochem, 114666), AZD4547 (Selleck, S2801), Chir99021 (Gift from D. Murphy), Cycloheximide (Sigma, 1810), EGF (Sigma, E4127), FGF2 (Thermo, PHG0263), GSK690693 (Gift from D. Murphy), PRN1371 (Selleck, S8578), Proteinase K (Thermo, 25530049), Pyr41 (Sigma, N2915), qVD-oph (AdooQ BioScience, A14915-25), Rapamycin (Santa Cruz, sc-3504), S55746 (ProbeChem, PC-63502), S63845 (Chemgood, C-1370), Sytox Green (Thermo, S7020) and Trametinib (Cambridge Bioscience, HY-10999).

### Lentiviral transduction

CRISPR/Cas9-based genome editing was performed using LentiCRISPRv2-puro (Addgene #52961) or LentiCRISPRv2-blasti ^9^ using the following guide sequences as previously described^38^: hBAX, 5’-AGTAGAAAAGGGCGACAACC-3’; hBAK, 5’-GCCATGCTGGTAGACGTGTA-3’; hERK1, 5’-CAGAATATGTGGCCACACGT-3’; hERK2, 5’-AGTAGGTCTGATGTTCGAAG-3’; hFGF2.1, 5’-TATGCAAGTCCAACGCACTG −3’ and hFGF2.2, 5’-CGAGCTACTTAGACGCACCC-3’.

For stable cell line generation, 5*10^6 293FT cells were transfected in 10 cm dishes using 4 μg polyethylenimine (PEI, Polysciences) per μg plasmid DNA with lentiCRISPR_V2 (Addgene 52961): gag/pol (Addgene 14887): pUVSVG (Addgene 8454) at a 4:2:1 ratio. After 48 and 72 hours of transfection, virus containing supernatant was filtered, supplemented with 1 μg/ml polybrene (Sigma) and added to 50.000 recipient cells in a 6 well plate. Selection with appropriate antibiotics (1 μg/ml puromycin (Sigma) or 10 μg/ml blasticidin (InvivoGen)) was started 24 h after the last infection and continued for one week.

### Supernatant assays

Generally, cells were treated for 3 h with 500 nM venetoclax, then the medium was replaced with regular growth medium. After an additional 45 h the supernatant was harvested, filtered and added onto recipient cells. For Proteinase K treatment, supernatant was treated with 200 μg/ml Proteinase K for 60 min at 50°C, followed by 5 min at 95°C. After cooling down, the treated supernatant was added to recipient cells. For centrifugal filtration, supernatant was filtered and added into an Amicon Ultra 15 ml tube (Merck) with a 3 kDa cut-off. After spinning at 4000 rpm for 60 min, the concentrate was diluted with regular growth medium to its original volume and the concentrate or the flowthrough was added onto recipient cells. The FGF2 ELISA was performed using the Human FGF-basic ELISA MAX Deluxe (Biolegend) according to the manufacturer’s instructions after 50x concentration of the supernatant by centrifugal filtration (see above). The final FGF2 concentration was determined using a standard of recombinant FGF2, taking into account the concentration step.

### Plasmid and siRNA transfection

For plasmid transfection, Lipofectamine 2000 was used according to the manufacturer’s instructions. Transfection of siRNA was performed using Oligofectamine according to the manufacturer’s instructions. The following siGENOME SMARTpool siRNAs from Dharmacon were used: Non-targeting, D0012061305; hFGFR1, M-003131-03-0005; hFGFR2, M-003132-04-0005; hFGFR3, M-003133-01-0005 and hFGFR4, M-003134-02-0005.

### Western blotting

Cell lysates were prepared using lysis buffer (1 % NP-40, 0.1 % SDS, 1mM EDTA, 150mM NaCl, 50mM Tris pH7.5, supplemented with Complete Protease Inhibitors (Roche) and PhosSTOP (Roche)). Protein concentration was determined with Bio-Rad Protein Assay Dye (5000006), and lysates were separated by SDS-PAGE followed by blotting onto nitrocellulose membranes and incubation with primary antibody (1:1000 in 5% milk) overnight. After washing in TBS/T and incubation with secondary antibody (Li-Cor IRDye 800CW donkey anti-rabbit, #926-32213), blots were developed on a Li-Cor Odyssey CLx system and acquired using Imagestudio (Li-Cor). The following primary antibodies were used: BAK (12105, Cell Signaling), BAX (2772, Cell Signaling), BCL-2 (2762, Cell Signaling), ERK1/2 (4695, Cell Signaling), FGFR1 (9740, Cell Signaling), FGFR3 (4574, Cell Signaling), FGFR4 (8562, Cell Signaling), GFP (In house), HSP60 (4870, Cell Signaling), MCL1 (5453, Cell Signaling), pERK1/2 (4370, Cell Signaling), CASPASE 3 (9662, Cell Signaling), CASPASE 9 (9502, Cell Signaling), PARP1 (9532, Cell Signaling) and cleaved CASPASE 3 (9664, Cell Signaling).

### Quantitative RT-PCR

RNA from cultured cells was isolated with the GeneJET RNA purification kit according to the manufacturer’s instructions. cDNA synthesis was performed according to the manufacturer’s instructions using the High Capacity cDNA Reverse Transcription Kit (Thermo), and qPCR was performed with the Brilliant III Ultra-Fast SYBR Green qPCR Master Mix (Agilent Technologies) on a QuantStudio 3 machine (Applied Biosystems) with the following program: 3 min at 95°C, 40 cycles of 20 s at 95°C, 30 s at 57°C, 30 s at 72°C and a final 5 min at 72°C. Results were analysed using the 2^−ΔΔCt^ method. The following primers were used: hACTIN s: 5’-CACTGTCGAGTCGCGTCC-3’, hACTIN a: 5’-GTCATCCATGGCGAACTGGT-3’,hBCL-2 s: 5’-CTGCACCTGACGCCCTTCACC-3’, hBCL-2 a: 5’-CACATGACCCCACCGAACTCAAAGA-3’, hCDX2 s: 5’-GACGTGAGCATGTACCCTAGC-3’, hCDX2 a: 5’-GCGTAGCCATTCCAGTCCT-3’,hDUSP6 s: 5’-GAAATGGCGATCAGCAAGACG-3’, hDUSP6 a: 5’-CGACGACTCGTATAGCTCCTG-3’,hMCL1 s: 5’-AAAGCCTGTCTGCCAAAT-3’, hMCL1 a: 5’-CCTATAAACCCACCACTC-3’ hSPRY4 s: 5’-TCTGACCAACGGCTCTTAGAC-3’ and hSPRY4 a 5’-GTGCCATAGTTGACCAGAGTC-3’.

### Cell death assays

Short term cell death was determined with an Incucyte FLR imaging system (Essen Bioscience) as previously described ^39^. Cells were treated as described in the Figure legend together with 30 nM Sytox green and imaged every 1 or 2 hours. Analysis was performed using the Incucyte software and the number of dead (Sytox green positive) cells was normalised to the confluency at t = 0. Long-term colony formation assay was performed by plating 1000 cells per 6 well and treatment as described in the Figure legend. After 48 h of treatment with supernatant, the medium was changed to 2.5 μM venetoclax/S63845. Medium was changed to regular growth medium after an additional 48 h, and resulting colonies were stained with crystal violet after an additional 5 days. Cell death analysis via FACS was performed as previously described ^40^. In short, treated cells were harvested and stained with 5 μg/ml propidium iodide and Annexin V (Biolegend) in Annexin V-binding buffer (10 mM Hepes pH 7.4, 140 mM NaCl, 2.5 mM CaCl_2_) for 15 minutes. Flow cytometry was conducted on a BD FACSCalibur machine; cells negative for propidium iodide and Annexin V were considered alive.

### Membrane Ligand array

Supernatant was harvested from HeLa tBID2A cells as described above and used to probe a Human Growth Factor Antibody Array (Abcam, ab134002) according to the manufacturer’s instructions.

### Bioinformatic analysis

Mutation, survival and gene expression data (Thymoma, Uterine Corpus Endometrial Carcinoma and Kidney Renal Clear Cell Carcinoma) was downloaded from cBioPortal (www.cbioportal.org). Samples with alterations in components of the FGFR signalling pathway (FGFR1, FGFR3, GRB2, FRS2, SOS1, HRAS, RAF1, MAPK1, MAPK3) were removed. The FGF score was determined by averaging the expression of the fgf2 induced gene set (CDH2, CDX4, FES, FGF2, FRS2, FYN, HOXA10, INHBA, MAP2K1, MAPK8, MMP7, PF4, RUNX2, SERPINF1) from the Harmonizome database (http://amp.pharm.mssm.edu/Harmonizome/). Pearson correlation was calculated between FGF score and BCL-2 or MCL1 expression. Samples were stratified into thirds based on FGF score, BCL-2 or MCL1 expression. Survival was analysed comparing samples with high FGF score and high expression of BCL-2 or MCL1 with low FGF score and low expression of BCL-2 and MCL1. Analysis was conducted in R, the used code is available upon request.

## Acknowledgements

This work was supported by Cancer Research UK grant C40872/A2014 (S.W.G.T) and a Tenovus small pilot grant (F.J.B). We thank K. Campbell for reviewing the manuscript and all members of the Tait laboratory for helpful suggestions. We would like to thank the Core Services and Advanced Technologies at the Cancer Research UK Beatson Institute (C596/A17196), with particular thanks to the Beatson Advanced Imaging Resource, Biological Services Unit, Histology and Molecular Technologies. Figures were created with BioRender.com.

## Author Contributions

Conceived the study and designed the work plan: F.J.B and S.W.G.T; Experimental work: F.J.B, C.C, D.Z; Data analysis: F.J.B and S.W.G.T; Manuscript writing: F.J.B and S.W.G.T.

---

**Supplementary Figure 1:**
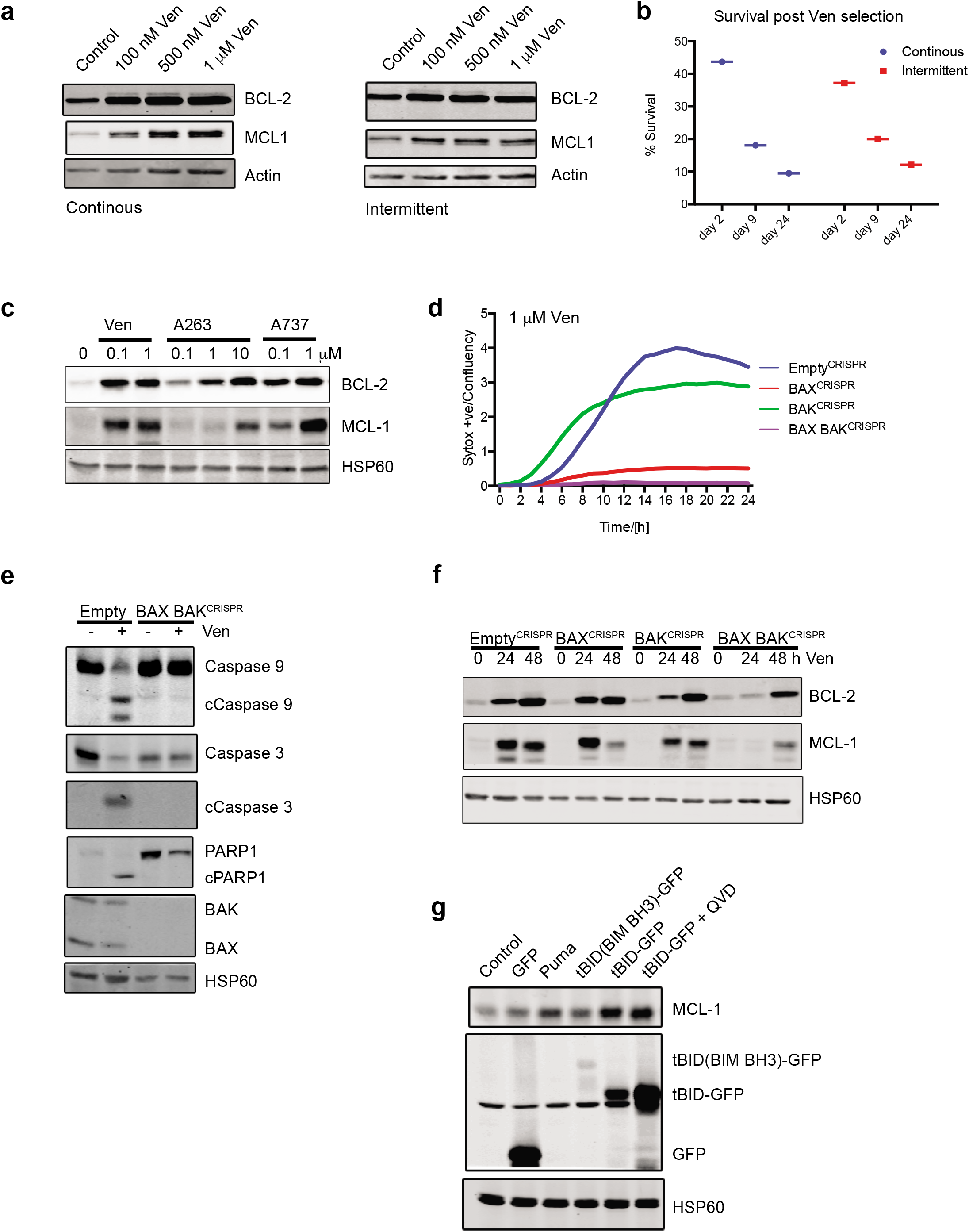
BH3-mimetics and BH3-only proteins upregulate BCL-2 and MCL1 causing apoptotic resistance. (a) HeLa tBID2A cells were continuously (left) or intermittently (right) cultured in different doses of venetoclax for 14 days, after which cells were harvested and protein expression determined by western blot. Images from one experiment (b) HeLa tBID2A cells made resistant as described in (a) were analysed for cell death by Annexin V/propidium iodide in response to venetoclax. The cells were cultured in regular medium for the indicated times, and apoptosis was determined by flow cytometry. Data from one experiment (c) HeLa tBID2A cells were treated for 48 h with the indicated BH3 mimetics, harvested and analysed for protein expression by western blot. Representative images from two independent experiments (d) Control, BAX^CRISPR^, BAK^CRISPR^ and BAX BAK^CRISPR^ HeLa tBID2A cells were treated with 1 μM venetoclax and cell viability was monitored by Sytox green staining and Incucyte live cell imaging. Representative images from two independent experiments (e) Control and BAX BAK^CRISPR^ HeLa tBID2A cells were treated with 500 nM venetoclax for 3 h, harvested and analysed for protein expression by western blot. Images from one experiment (f) Control, BAX^CRISPR^, BAK^CRISPR^ and BAX BAK^CRISPR^ HeLa tBID2A cells were treated with 500 nM venetoclax, harvested at the indicated time points and protein expression analysed by western blot. Representative images from two independent experiments (g) HeLa cells were transfected with the indicated constructs for 48 h and protein expression was analysed by western blot. Representative image one experiments

**Supplementary Figure 2:**
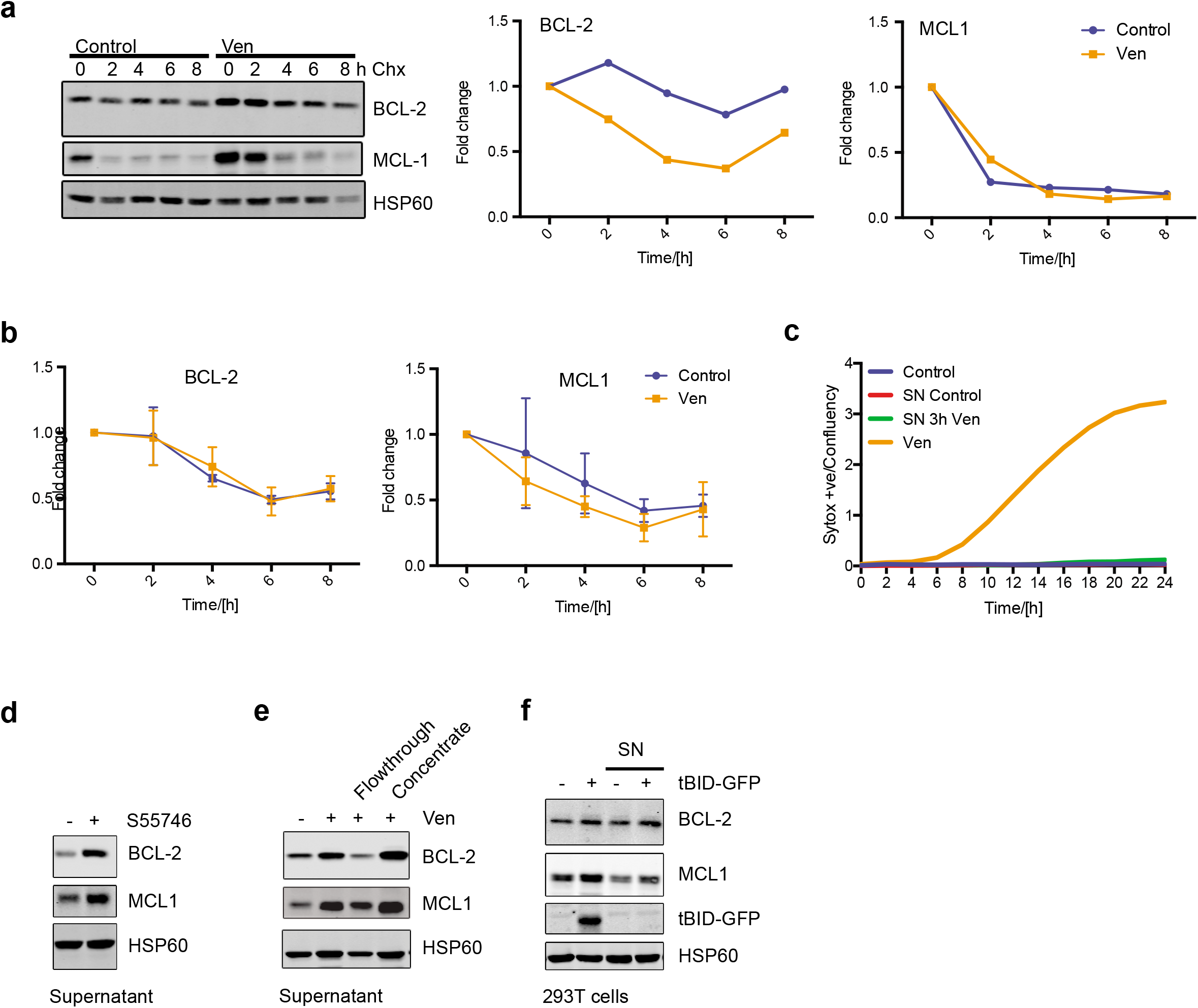
BH3-mimetics and BH3-only proteins can upregulate anti-apoptotic BCL-2 proteins in a non-cell autonomous manner. (a) HeLa tBID2A cells were treated with 500 nM venetoclax for 24 h, followed by treatment with 1 μg/ml cycloheximide. Cells were harvested at the indicated times and protein expression analysed by western blot. Quantification normalised to HSP60 relative to t = 0 shown on the right. Representative images from three independent experiments (b) HeLa tBID2A cells were treated with 500 nM venetoclax for 24 h, followed by treatment with 1 μM actinomycin D. Cells were harvested for the indicated times and RNA expression analysed by RT-qPCR. n = 3 independent experiments; mean values ± s.d. (c) HeLa tBID2A cells were treated as indicated and viability monitored by Sytox green uptake and Incucyte live cell imaging. Representative data from three independent experiments (d) Supernatant from control or HeLa tBID2A cells treated with 100 nM S55747 was added to recipient cells and protein expression was analysed by western blot after 48 h. Representative images from two independent experiments (e) Supernatant from control or venetoclax treated HeLa tBID2A cells was harvested and filtered with a 3 kDa cut-off centrifugal spin column. The concentrate was subsequently adjusted to its original volume and flowthrough and concentrate was added onto recipient cells. After 48 h, cells were harvested and protein expression analysed by western blot. Representative images from two independent experiments (f) 293T cells were transfected with 1 μg tBID-GFP for 24 h or treated with supernatant from tBID-GFP transfected 293T cells. After 24 h, the cells were harvested and protein expression was analysed by western blot. Representative images from two independent experiments

**Supplementary Figure 3:**
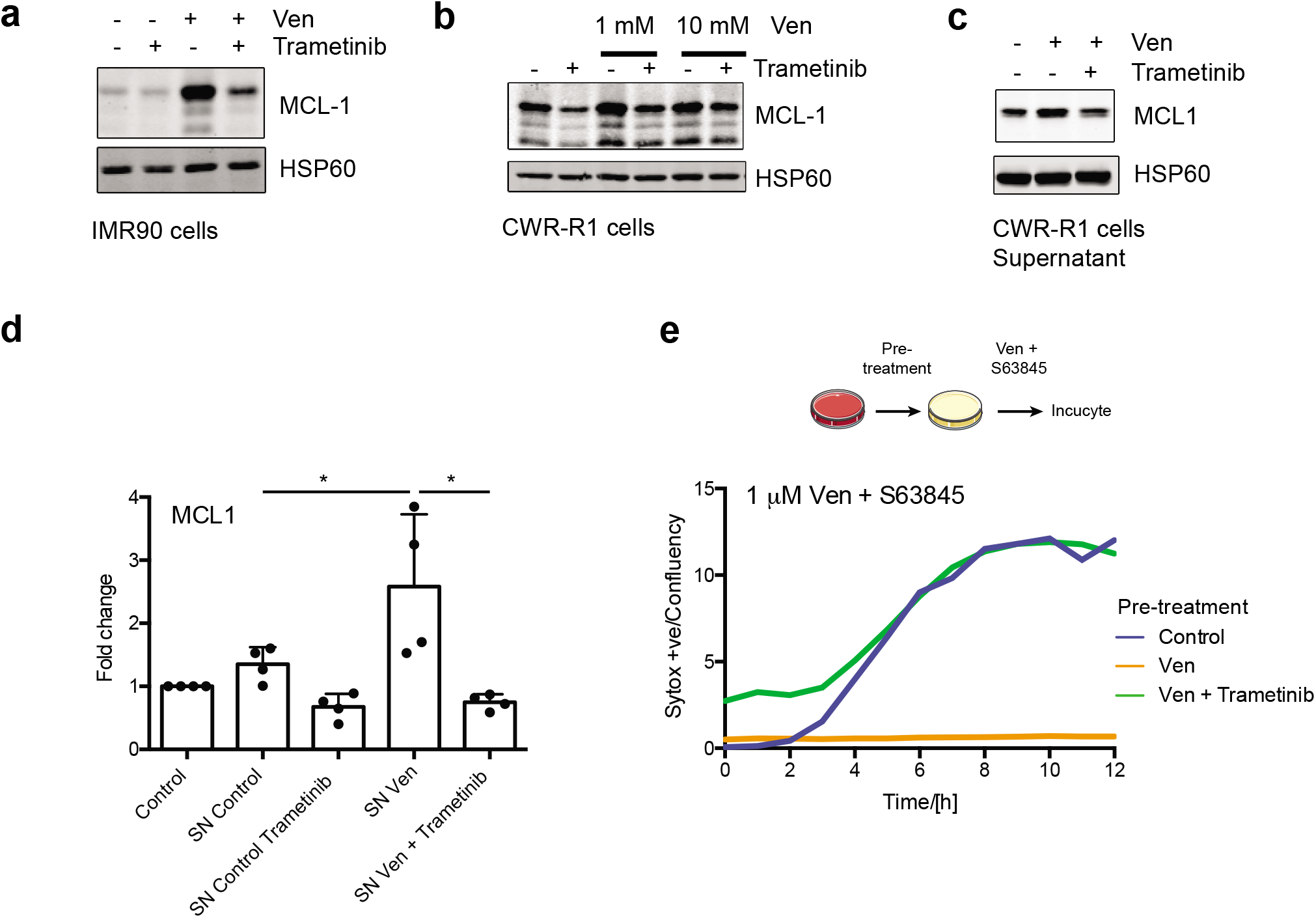
Non-cell autonomous upregulation of anti-apoptotic BCL-2 proteins requires MEK/ERK signalling. (a) IMR90 cells were treated with 10 μM venetoclax in combination with trametinib as indicated. After 48 h, cells were harvested and protein expression analysed by western blot. Representative images from two independent experiments (b) CWR-R1 cells were treated with venetoclax in combination with trametinib as indicated. After 48 h, cells were harvested and protein expression analysed by western blot. Representative images from two independent experiments (c) Supernatant from control or venetoclax treated CWR-R1 cells was treated with 500 nM trametinib as indicated before addition onto recipient cells and protein expression was analysed by western blot after 48 h. Representative images from two independent experiments (d) Supernatant from control or venetoclax treated HeLa tBID2A cells was treated with 500 nM Trametinib as indicated before addition onto recipient cells and RNA expression was analysed by RT-qPCR after 48 h. n = 4 independent experiments; mean values ± s.d., *: p < 0.05, Tukey corrected one-way ANOVA (e) HeLa tBID2A cells were treated with or without venetoclax in combination with 500 nM Trametinib as indicated for 48 h followed by treatment with venetoclax and S63845. Cell viability was then monitored by Sytox Green staining and Incucyte imaging. Representative data from two independent experiments

**Supplementary Figure 4:**
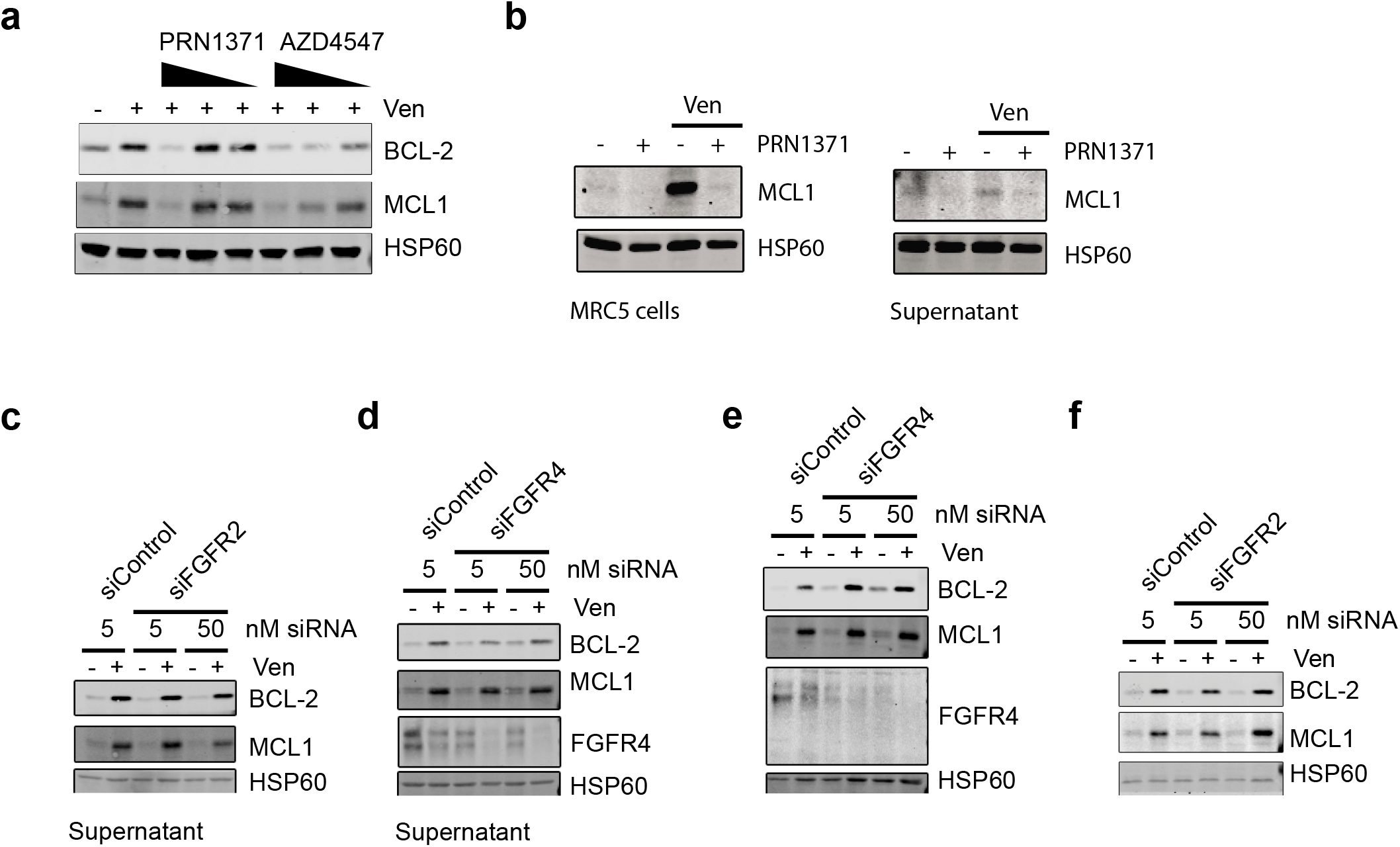
FGF signalling mediates non-cell autonomous upregulation of BCL-2 proteins. (a) HeLa tBID2A cells were treated with venetoclax in combination with decreasing doses of FGFR inhibitors as indicated, cells were harvested after 48 h and protein expression was analysed by western blot. Representative images from tree independent experiments (b) MRC5 cells were treated with venetoclax in combination with 10 μM PRN1371 (left) or with supernatant from MRC5 cells treated with venetoclax and addition of 10 μM PRN1371 (right) as indicated for 48 h, after which cells were harvested and protein expression was analysed by western blot. Representative images from two independent experiments (c) – (f) HeLa tBID2A cells were transfected with the indicated amount of siRNA targeting FGFR2 or FGFR4 for 24 h before addition of supernatant from control cells treated with 500 nM venetoclax (c, d) or direct treatment with 500 nM venetoclax (e, f). Cells were harvested after 48 h and protein expression was analysed by western blot. Representative images from three independent experiments

**Supplementary Figure 5:**
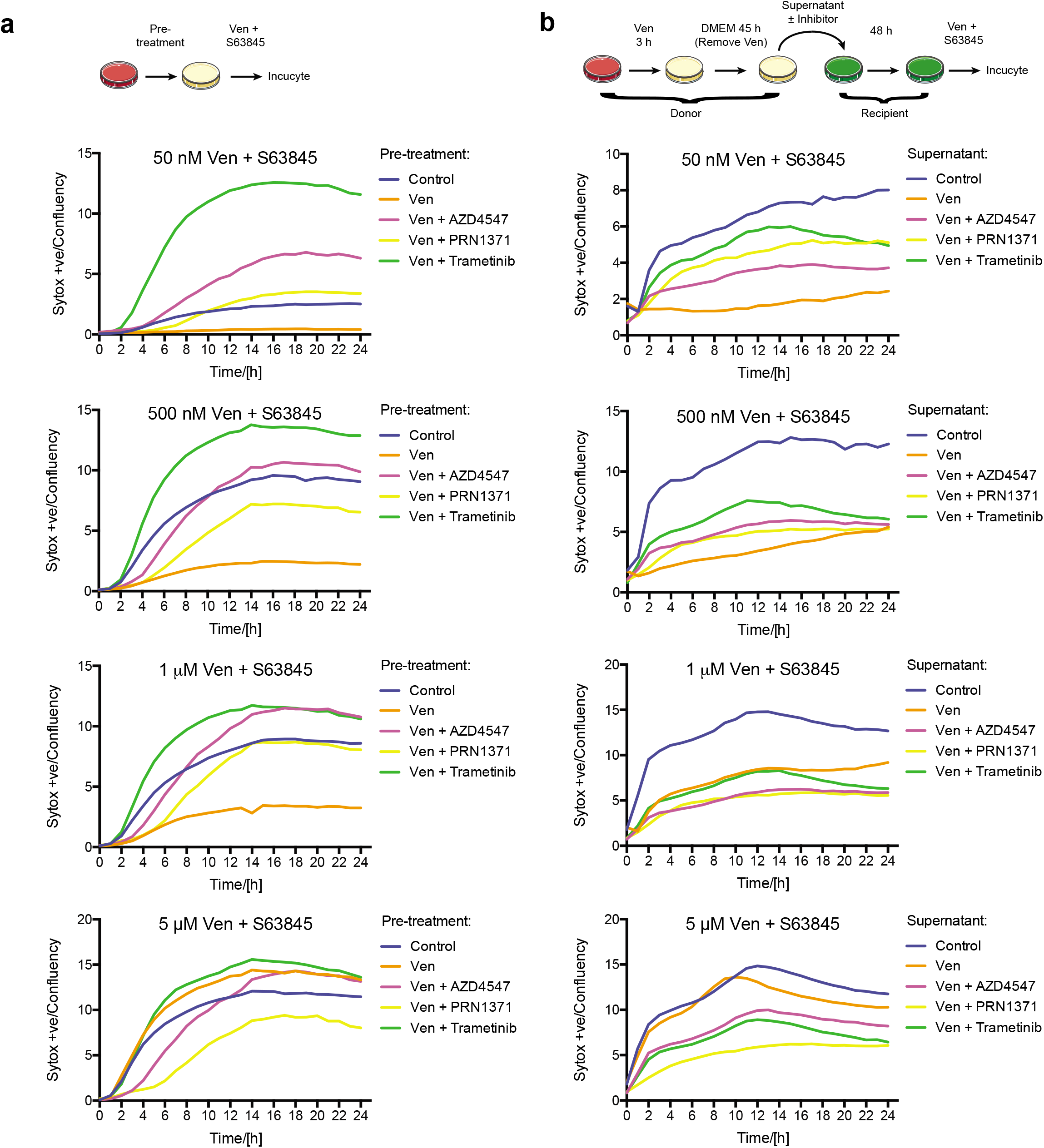
FGF signalling is essential for non-cell autonomous apoptotic resistance. (a) HeLa tBID2A cells were treated with 500 nM venetoclax in combination with RTK pathway inhibitors (AZD4547 (5 μM), PRN1371 (10 μM) or Trametinib (500 nM)) as indicated for 48 h. Then the cells were treated with venetoclax + S63845 and cell survival was monitored by Incucyte. Representative images from two independent experiments (b) HeLa tBID2A cells were incubated with 500 nM venetoclax treated supernatant supplemented with RTK pathway inhibitors (AZD4547 (5 μM), PRN1371 (10 μM) or Trametinib (500 nM)) as indicated before addition onto recipient cells for 48h. Then the cells were treated with venetoclax + S63845 and cell survival was monitored by Incucyte. Representative images from two independent experiments

**Supplementary Figure 6:**
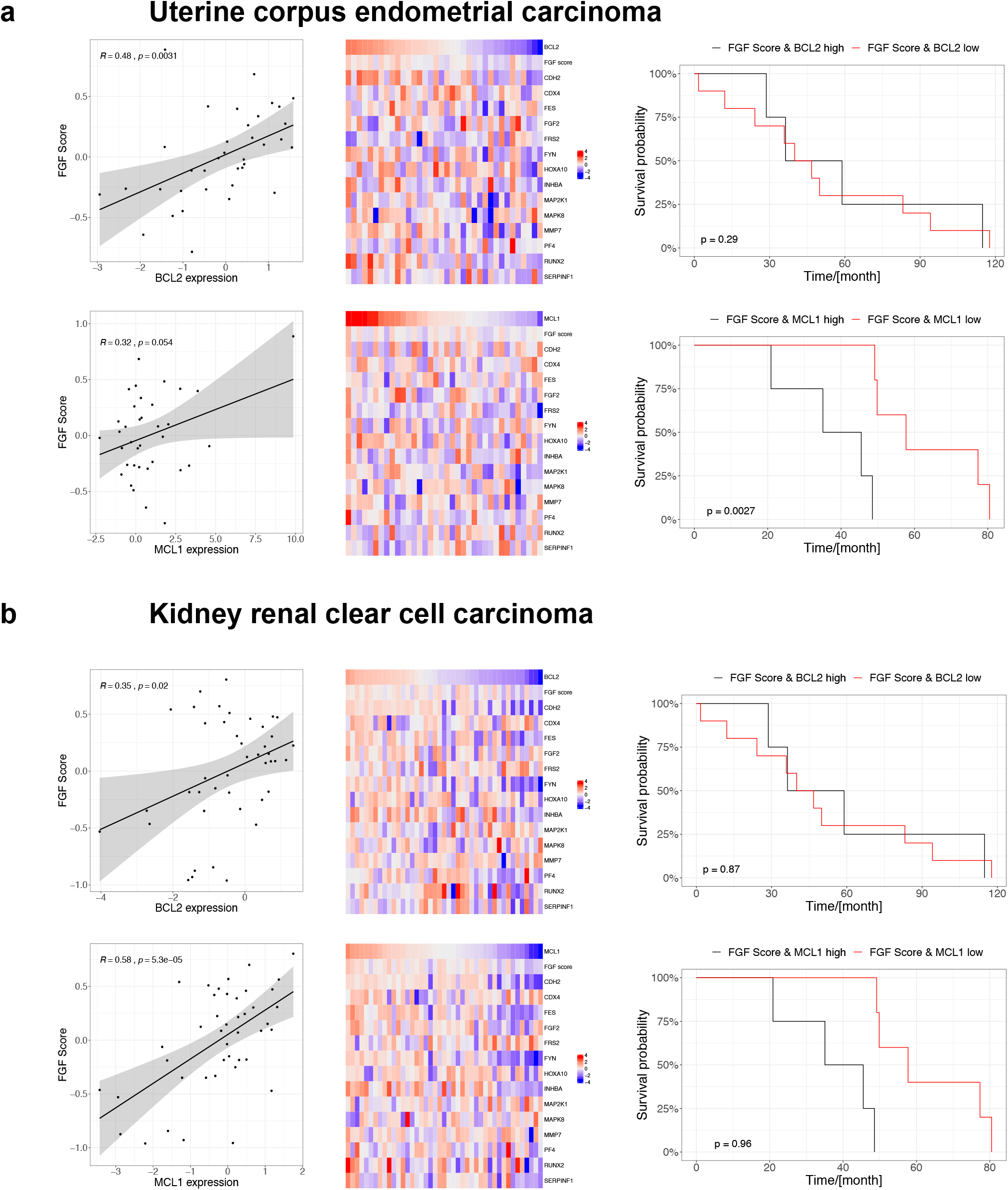
Correlation between FGF signalling and BCL2 family member expression predict patient survival in some cancer types. (a) Correlation between FGF Score and BCL-2 (upper left) or MCL1 (lower left) in the TCGA Uterine corpus endometrial carcinoma dataset. FGF score, FGF receptor target gene and BCL2 (upper middle) or MCL1 (lower middle) expression in the TCGA Uterine corpus endometrial carcinoma dataset. Survival of TCGA Uterine corpus endometrial carcinoma patients stratified by high FGF score and BCL2 (upper right) and MCL1 (lower right) expression (b) Correlation between FGF Score and BCL-2 (upper left) or MCL1 (lower left) in the Kidney renal clear cell carcinoma dataset. FGF score, FGF receptor target gene and BCL2 (upper middle) or MCL1 expression (lower middle) in the TCGA Kidney renal clear cell carcinoma. Survival of TCGA Kidney renal clear cell carcinoma patients stratified by high FGF score and BCL2 (upper right) and MCL1 (lower right) expression

